# Mechanical and possible auxetic properties of human Achilles tendon during in vitro testing to failure

**DOI:** 10.1101/2021.09.09.459526

**Authors:** Christopher V. Nagelli, Alexander Hooke, Nicholas Quirk, Consuelo Lopez De Padilla, Timothy E. Hewett, Martijn van Griensven, Michael Coenen, Lawrence Berglund, Christopher H. Evans, Sebastian A. Müller

**Author notes:** Address for correspondence: C. H. Evans, PhD, Rehabilitation Medicine Research Center, Mayo Clinic, 200, First Street SW, Rochester, MN 55905.

## Abstract

The Achilles tendon is the strongest tendon in the human body, but the basis of its high tensile strength has not been elucidated in detail. Here we have loaded healthy, human, Achilles tendons to failure in an anatomically authentic fashion while studying the local three-dimensional deformation and strains in real time, with very high precision, using digital image correlation (DIC). These studies identified a remarkable degree of anisotropic, medio-lateral auxetic behavior, with Poisson’s ratios not exceeding minus 1 in any part of the tendon at any time; under certain loads, discrete areas within the tendon had a Poisson’s ratio below minus 6. Early in the loading cycle, the proximal region of the tendon accumulated high lateral strains while longitudinal strains remained low. This behavior shielded the mid-substance of the tendon, its weakest part, from high longitudinal strains until immediately before rupture. These new insights are of great relevance to understanding the material basis of tendon injuries, designing improved prosthetic replacements, and developing regenerative strategies.

## Introduction

Tendons connect skeletal muscles to bones, enabling movement by transmitting the forces generated by muscle contraction. They comprise highly ordered, hierarchical collagenous structures that withstand large loads when stretched. They are frequently injured during sporting activities and other trauma (Schepsis *et al*., 2002); for unknown reasons, the incidence of tendon ruptures has increased in recent years (Ganestam *et al*., 2016; Lantto *et al*., 2015). Because tendon injuries do not heal well (Docheva *et al*., 2015; Muller *et al*., 2015) and interfere with quality of life, they create an increasing medical, economic and social burden (Hopkins *et al*., 2016). A detailed understanding of the material properties of tendons, as well as strain patterns leading to rupture, promises to reduce the incidence of such injuries and provide important new information of value for developing better regenerative strategies and superior prosthetic replacements.

The Achilles tendon connects the calf muscles to the heel (calcaneus; Fig. 1). It is the largest and strongest tendon in the human body, and resists forces that exceed 12-times body weight during simple activities such as running (Komi *et al*., 1992). However, the basis for the unusual strength of the normal human Achilles tendon is poorly understood. It is often assumed that its strength reflects its large size, but such a simplistic explanation ignores the fact that the Achilles tendon narrows to a width of only 1.8 cm at its mid-substance (Del Buono *et al*., 2013) where most Achilles tendon ruptures occur.

**Figure 1:**
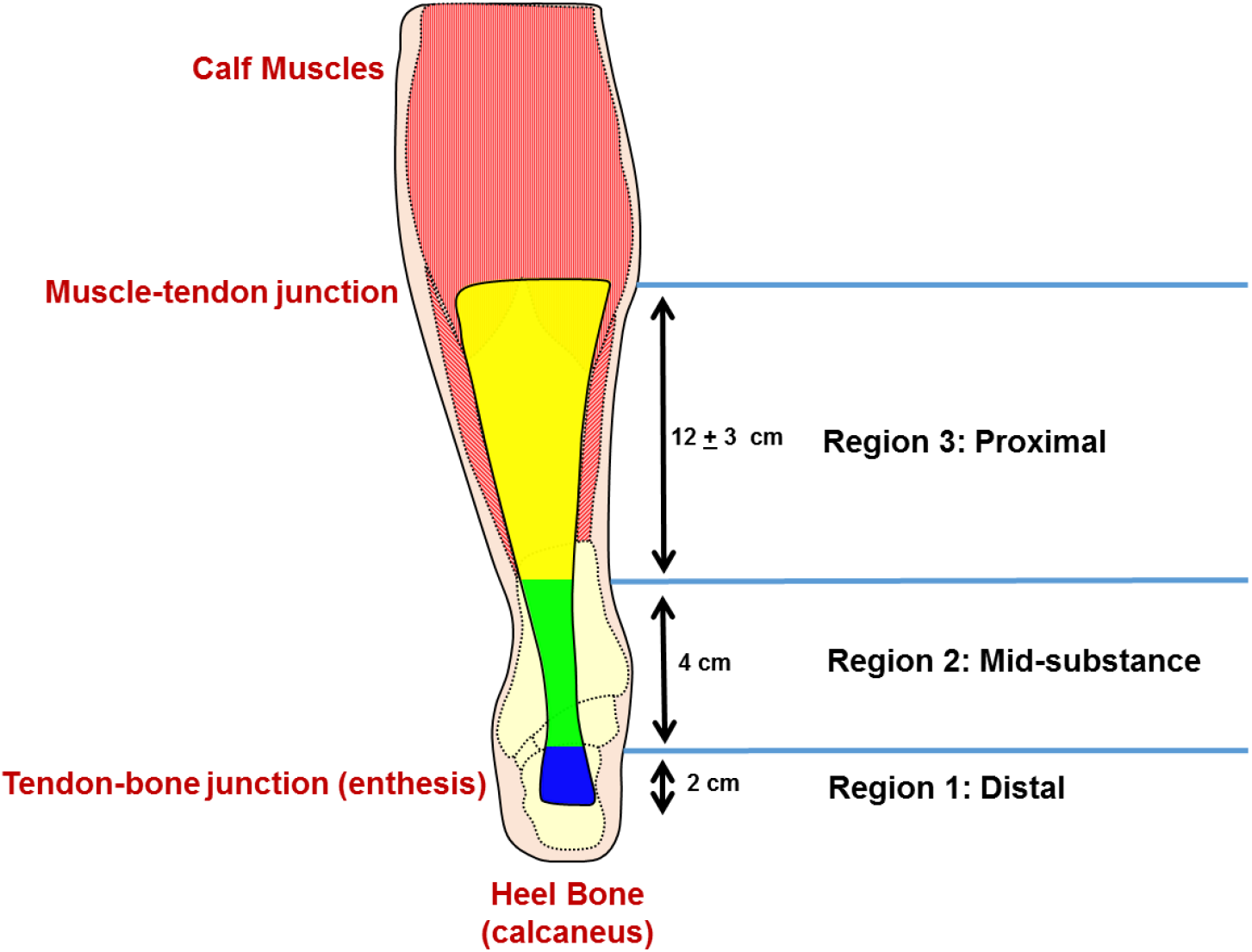
A schematic of the human Achilles tendon, showing division into 3 regions of analysis.

The lack of information concerning the material properties of the human Achilles tendon results partly from practical and technical challenges encountered when attempting to study this tissue *ex vivo*. The limited availability of normal, fresh, human Achilles tendons is one factor. Another challenge is the ability to fix Achilles tendon for testing in an anatomically authentic fashion, with its calcaneal insertion intact at one end and the calf muscles secured at the other exactly in line with the axis of loading. The magnitude of these constraints is reflected in the dearth of literature on the subject; we could identify only three published papers in which the material properties of freshly excised, normal, human Achilles tendons were measured during loading to failure (Louis-Ugbo *et al*., 2004; Wren *et al*., 2003; Wren *et al*., 2001); one additional study used Achilles tendon removed from embalmed cadavers (Lewis and Shaw, 1997). A particularly severe additional complication is the non-uniform cross section of the Achilles tendon, which renders the stresses and strains highly anisotropic along the length of the tendon.

For the study described here, our group obtained fresh, uninjured Achilles tendons from relatively young human donors. We then developed embedding techniques that retained the native distal and proximal insertions of the tendon in the calcaneus and calf muscles, respectively, while positioning the calcaneus-tendon-muscle unit in an anatomically correct alignment; this allowed mechanical testing to failure. In addition, digital image correlation (DIC) allowed us to measure, for the first time, local three-dimensional deformation and strains in real time with very high precision during loading to failure. Therefore, the main objective was to use the DIC technology to study the mechanical behavior of healthy Achilles tendons as they were failure tested under tension. We hypothesized that this group of young tendons would fail at exceeding high loads (>6,000 Newtons), and that we would gain insight into the deformation pattern as the tendons were stretched to failure. As well as demonstrating considerable regional and temporal strain heterogeneity, DIC revealed a remarkable degree of auxetic behavior. The overall average Poisson’s ratio never rose above -1 as the tendon was loaded to failure and in particular locations reached below - 6 under certain conditions.

## Materials and Methods

### Sample Preparation

Fresh frozen, complete human cadaveric lower extremities were obtained (Anatomy Gifts Registry, Hanover, MD) from 11 donors (6 male/ 5 female) aged 42.1 ± 9.8 years (height 1.78 ± 0.1 m; weight 83.3 ± 22.3 kg), with no history of traumatic lower extremity injury or surgery. The entire Achilles tendon, including the calcaneus and distal third of the *gastrocnemius* and *soleus* muscles, was harvested from each donor after which the enthesis was carefully exposed by dissection. A board certified orthopaedic surgeon (SAM) examined the tendons for macroscopic signs of injury or degeneration. Samples were secured in a metal fixture using bone cement and K-wire wrapping such that a portion of the calcaneus was above the surface with the enthesis clearly visible (Fig. 2). The samples were kept hydrated during dissection with saline.

**Figure 2:**
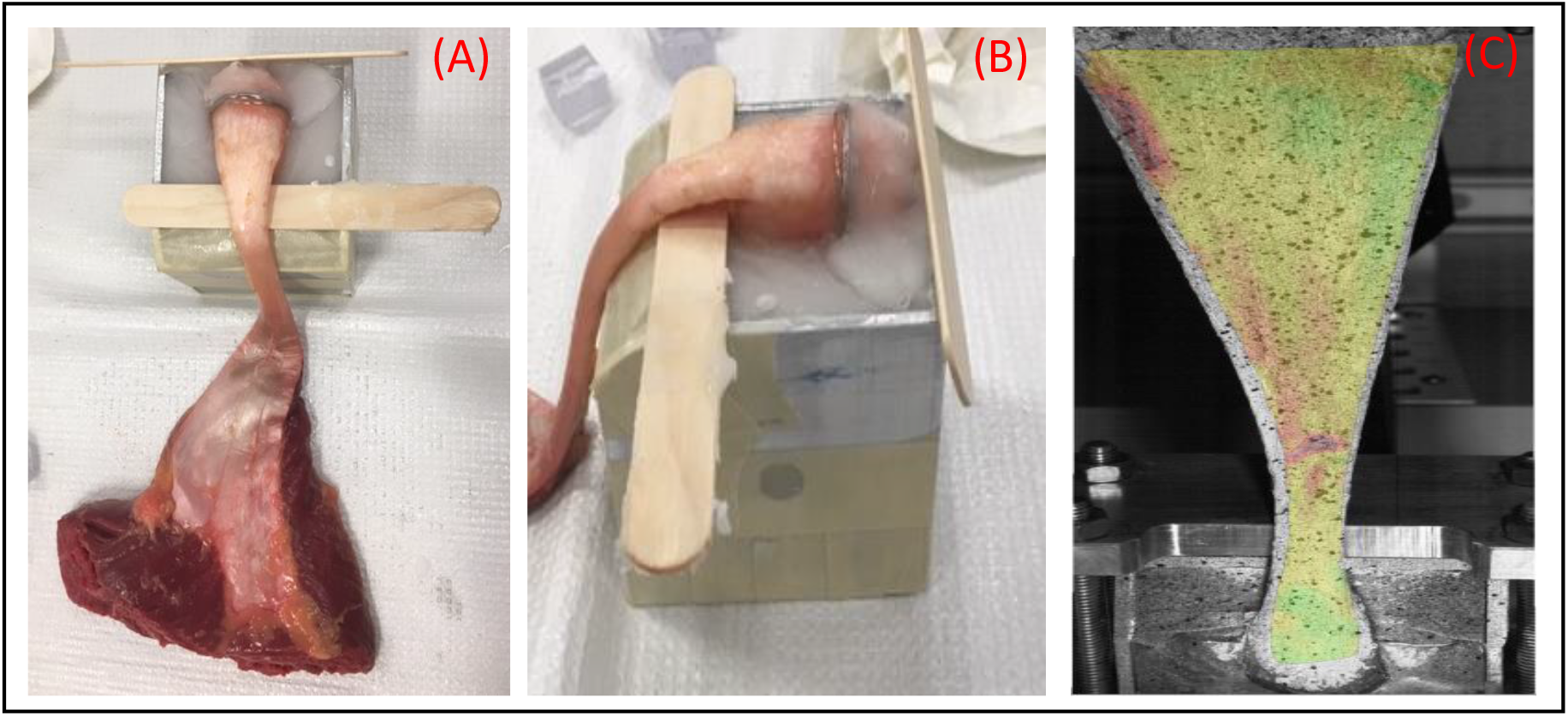
Frontal (A) and sagittal (B) plane views of the Achilles tendon insertion site with calcaneus exposed and secured with bone cement in a metal block. The experimental set-up (C) includes simultaneous tensile testing and local strain measurements using a digital imagine correlation system. The Achilles tendon is in an anatomically correct position with the calcaneus position at a 30º angle, to simulate the natural position of a plantigrade foot in a neutral position. The midsubstance of the tendon did not have contact with the frame during testing.

### Mechanical Testing

Samples were loaded into a servo-hydraulic testing machine (model 312, MTS System Corporation, Eden Prairie, MN, USA) using a customized fixator with the calcaneus at a 30°angle to simulate the anatomic position of a plantigrade foot in a neutral position. The muscle and musculotendinous junction were fixed with a custom cryo-clamp. Tendons were pre-conditioned under 10 cycles of pre-load to 2% strain and then subjected to displacement-controlled tensile testing at 5 mm/s until failure. Force was measured with a 2500kg load cell (model 3397; Lebow products, Troy MI; accuracy 0.05%). Force and displacement data were collected at 128 Hz.

### Digital Image Correlation

For DIC measurements, the posterior surface of each tendon was patterned first with a solid white, mineral-oil based face paint with black speckling applied on top of the white base layer using spray paint. The samples were kept hydrated during testing with saline which was carefully applied using a spray bottle not to distort the speckling pattern. DIC imaging was performed using a GOM-ARAMIS 4M system (GOM-Optical Measuring Techniques, Braunschweig, Germany). The system consisted of two cameras with a resolution of 4 megapixels (2400 × 864 pixels) equipped with 35 mm lenses. The tendon was illuminated by two LED lights (Schneider Optische Werke, GmbH, Bad Kreuznach, Germany). The DIC system was calibrated prior to the tests to accommodate a measurement volume of 235 × 170 × 160mm. Images were collected using the ARAMIS DIC software (ARAMIS v2018, GOM, Braunschweig, Germany) at a frame rate between 32 and 384 Hz with all specimens down sampled to 32 Hz during analysis. Image processing was performed using the ARAMIS DIC software using a subset size of 19 pixels, a step size of 12 pixels, and a virtual strain gauge length of 2.2 mm.

### Histology

Samples (average 15 mm in width) were dissected from areas proximal and distal to the rupture site, fixed in 10% neutral buffered formalin for 48 h, dehydrated through graded alcohols and embedded in paraffin. Sections (5 µm) were rehydrated and stained with hematoxylin-eosin or alcian blue, pH 2.5. Stained sections were examined microscopically and assigned a Bonar score, a validated measure of tendon degeneration based upon cellularity, tenocyte shape, collagen structure, vascularity and ground substance (Maffulli *et al*., 2008). This provides an aggregate score between 0 (normal) and 12 (severely abnormal). Bright field images were acquired using an automated inverted microscope (Olympus IX83, Waltham, MA). Images were stitched at 20X using cell Sense Olympus imaging software.

### Data Analysis

Primary outcome measurements were: failure load, stiffness, 3 components of engineering strain (longitudinal strain (superior-inferior direction), transverse strain (medio-lateral direction), and major strain), and Poisson’s ratio. Stiffness was computed from the linear region of the force displacement curve. This is the region of the curve between the toe region and the yield point. The ARAMIS DIC software calculates finite strains. Major strain is defined as strain in the direction of greatest deformation. This means the direction of the major strain can vary across the surface of a material. While initially developed in the sheet metal industry, the introduction of optical metrology as a tool in other fields such as biomechanics has expanded the value of major strain as a uniquely informative metric. Computationally, circular marks are digitally created using the speckling pattern on the tissue in its unloaded state. As the tissue is loaded into tension, these digital circles become ellipses whose long and short axes define major and minor strain, respectively, as indicated in figure 3. Each strain variable of interest and Poisson’s ratio were analyzed by both studying their heat maps and by averaging their values across the entire tendon surface and resampling them across a normalized load from the beginning of testing (0% of failure load) to failure (100% of failure load).

**Figure 3:**
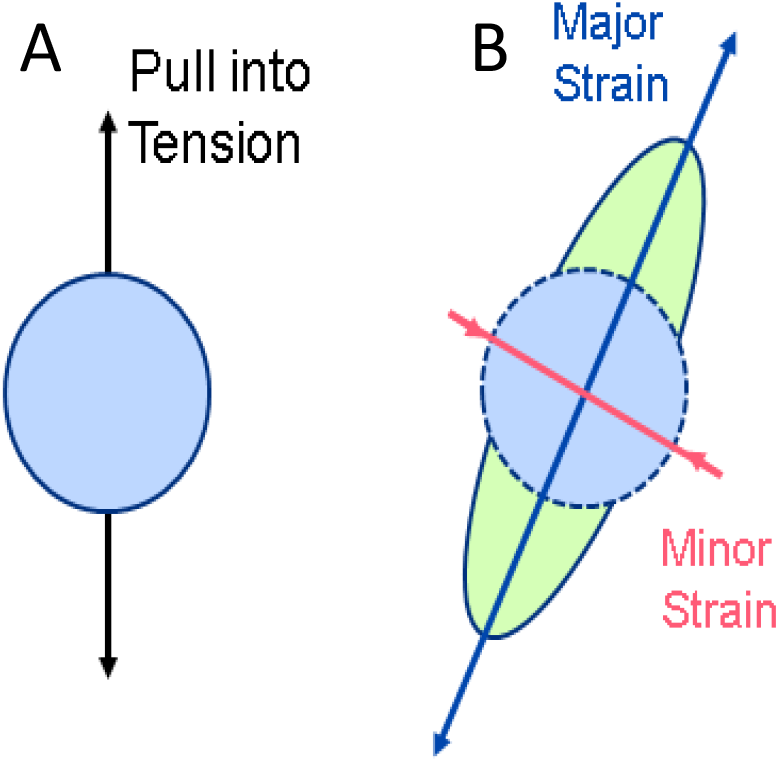
Detailed breakdown of major strain. Major strain is defined as the direction of highest deformation, and minor strain is defined as the direction of the lowest deformation. A circular mark placed is placed on the tendon prior to tensile testing (A). After testing, the post-testing ellipse provides an indication of major and minor strain (B).

To analyze more closely the regional differences in strain during mechanical testing, tendons were divided into three regions (Fig. 1): distal, calcaneal insertion site to 2 cm proximal (region 1); mid-substance, 2-6 cm proximal from calcaneal insertion (region 2); proximal, > 6 cm proximal from calcaneal insertion to musculotendinous region (region 3).

Analysis of variance (ANOVA) tests were performed to determine significant differences between failure types for failure load and stiffness (α=0.05).

## Results

### Mechanical testing

Five tendons failed in the mid-substance (region 2; figure 1) and five underwent calcaneus avulsions. One sample failed via muscle tear before the tendon ruptured and was not included when performing ANOVA analysis of the mechanical testing data. Mid-substance failures occurred at an average failure load of 5,649.1 ± 565.5 N, while calcaneus avulsion failures occurred at a failure load of 4,216.9 ± 1,196.4 N (Fig. 4A; p=0.04). Tendons that underwent calcaneus avulsions had significantly lower average tendon stiffness than those failing in the mid-substance failures (Fig. 4B; p=0.002). There was no difference in the mechanical properties of tendons from male and female donors.

**Figure 4:**
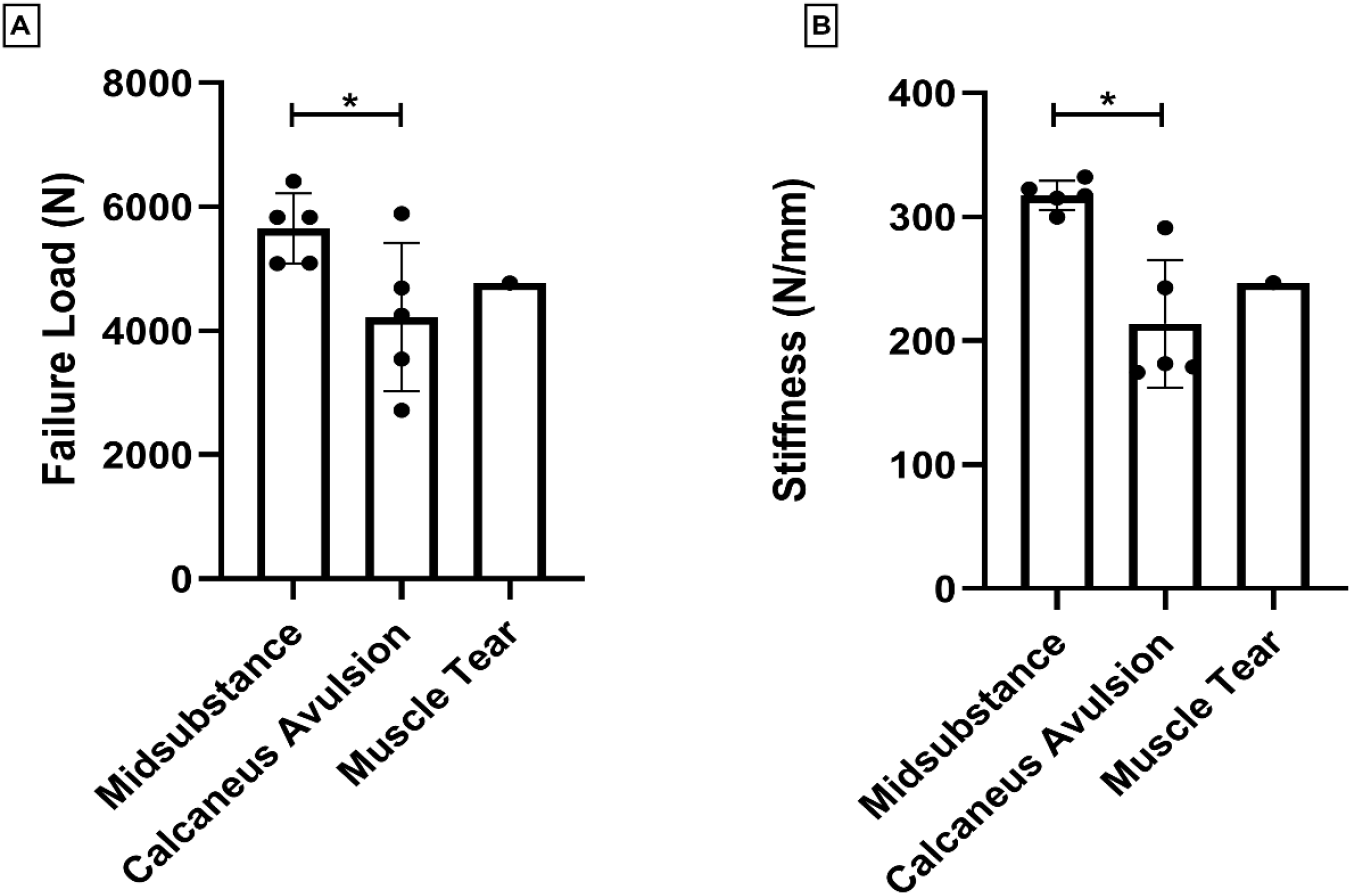
Differences in failure load (A) and tendon stiffness (B) between the tendon failure types. *Significant differences (p<0.05) were found between the mid-substance failures and calcaneus avulsion failures for failure load and tendon stiffness.

### Histology

Examination of the histological specimens provided Bonar scores of 0 - 0.5, in agreement with the gross anatomy observations indicating that the tendons were without preexisting tendinopathy. Representative histological images are shown in figure 5.

**Figure 5:**
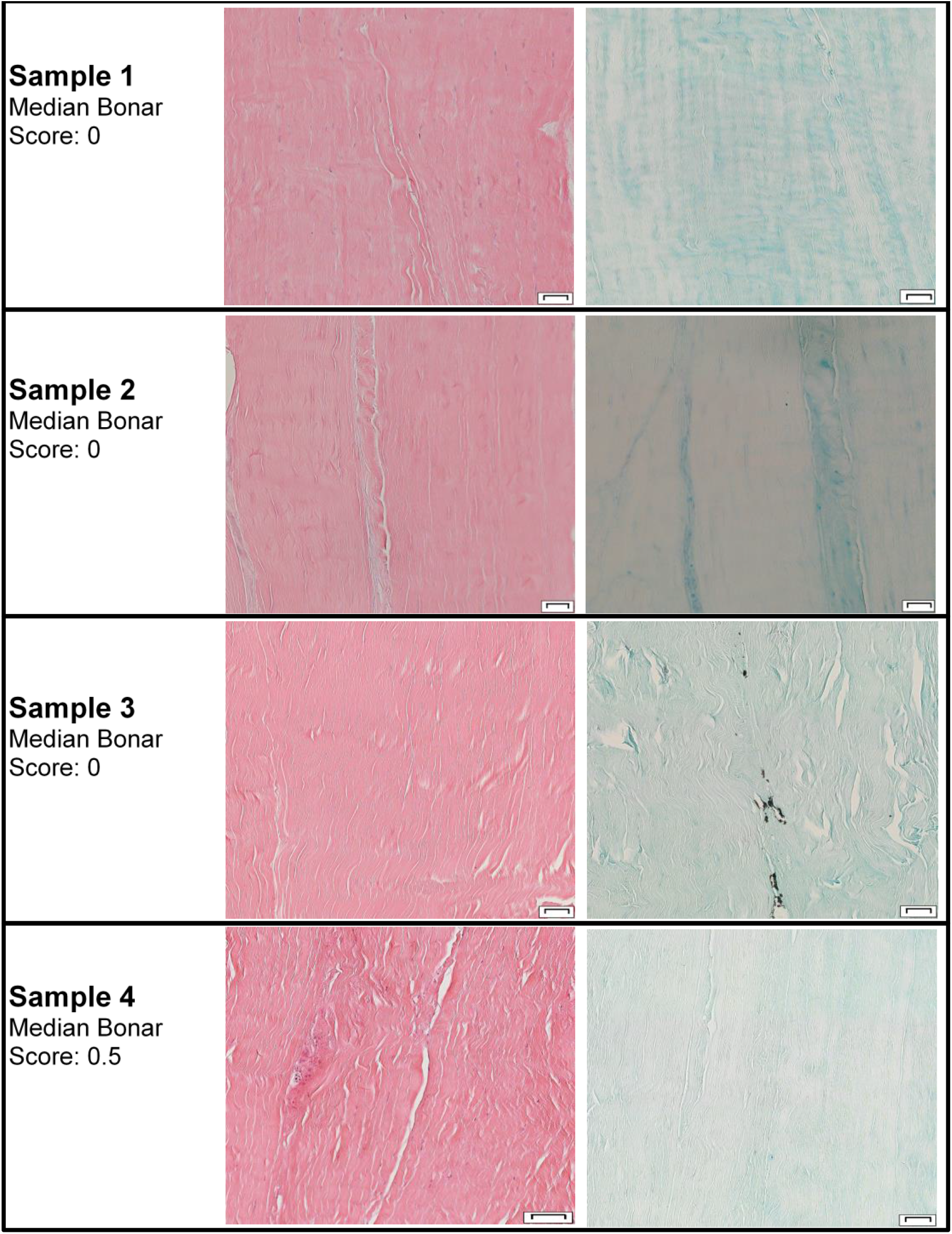
Representative histological sections of tendons stained with hematoxylin and eosin or alcian blue. Of note, histological images appear normal with no indication of prior disease of the tendon or breakdown of structural integrity. No specimen had a median Bonar score > 0.5. Scale bars represent 50 µm in length.

### Strain Analysis

All eleven samples were included in the DIC assessment. Table 1 includes transverse and longitudinal strains at a percentage of failure load (25%, 50%, 75%, and 100%) from each region for the tested Achilles tendon samples. As shown in the accompanying video (supplemental data), the deformation behavior varied considerably from region to region within a tendon, and also shifted dramatically as loading continued.

Early in the loading cycle considerable transverse strain accumulated in the proximal region (region 3, figure 1) of the tendon, with little longitudinal or transverse strain apparent elsewhere. However, shortly before failure, significant longitudinal strain appeared for the first time in the location that would go on to rupture. Figure 5 shows the directional breakdown of strain in a representative Achilles tendon sample at the moment it sustained a mid-substance failure. The heat map of major strain (Fig. 6A) shows hotspots near the calcaneus region, at the site of tendon failure and proximally around the musculotendinous region. Upon further analysis of the major strain’s directional component (Fig. 6B), it was clear that the direction of major strain varied widely across the surface of the tendon. Just prior to rupture, the more distal hotspots close to the failure site demonstrated major strain in the longitudinal direction for the first time, parallel to the direction of loading, while in the more proximal region (region 3, figure 1) the major strain continued to be in the transverse direction, orthogonal to the loading direction.

**Figure 6:**
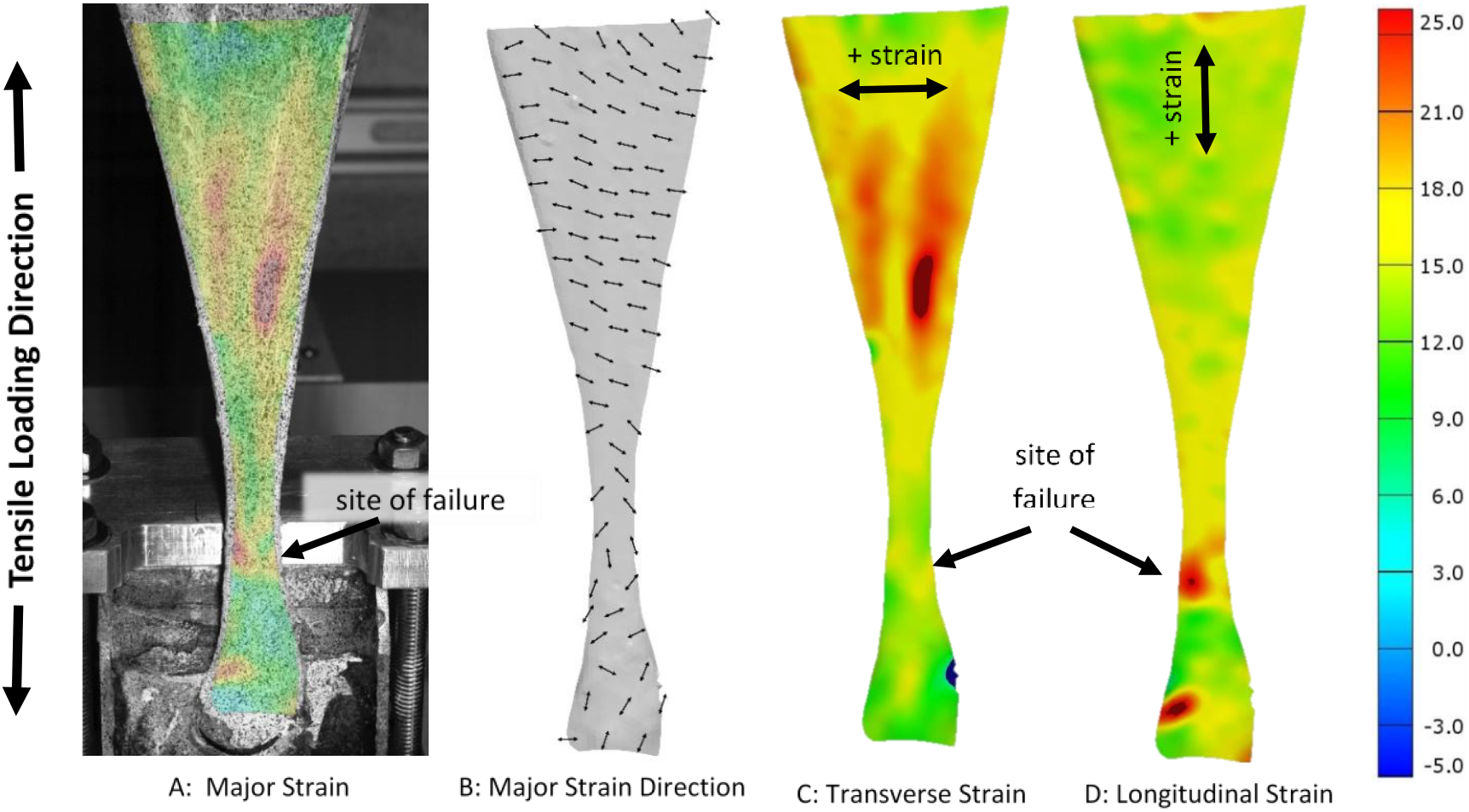
Directional breakdown of technical strain in one representative sample of a human Achilles tendon at rupture. The direction of strain as measured by major strain (A) and its directionality (B) indicates differences in strain direction at the site of rupture. In addition, at rupture, the tendons experienced greater transverse strain (C) than longitudinal strain (D), in the proximal region (region 3), whereas the opposite is true at the rupture site. (Supplemental video shows evolution of strain in real time during loading to failure).

This observation was confirmed when the heat maps for the transverse strain (Fig. 6C) and longitudinal strain (Fig. 6D) were compared. It was striking how the tendons experienced transverse strain and that localized regional differences in strain patterns clearly existed. A strikingly consistent strain pattern was seen for all specimens, regardless of whether they failed in the mid-substance or by calcaneus avulsion. The pooled data are represented in figure 6, which shows the average longitudinal and transverse strains as a function of percent failure load for the entire surface and for regions 1, 2, and 3.

Overall, transverse strain increased dramatically as loading increased, with a plateau at an average value of 9.5% strain. Average longitudinal strain increased more slowly, did not plateau and reached 6% strain at the time of rupture (Fig. 7A). However, there were striking regional differences. Thus, in regions 2 (mid-substance) and 3 (proximal), transverse strain exceeded longitudinal strain throughout almost the entire tensile test, especially at low and medium loads (Fig. 7C, D). As loading increased, the transverse strain eventually plateaued, while longitudinal strain increased rapidly until failure. In region 1 (adjacent to calcaneal insertion) the opposite was true, and the longitudinal strain exceeded transverse strain at all but the lowest loads (Fig. 7B).

**Figure 7:**
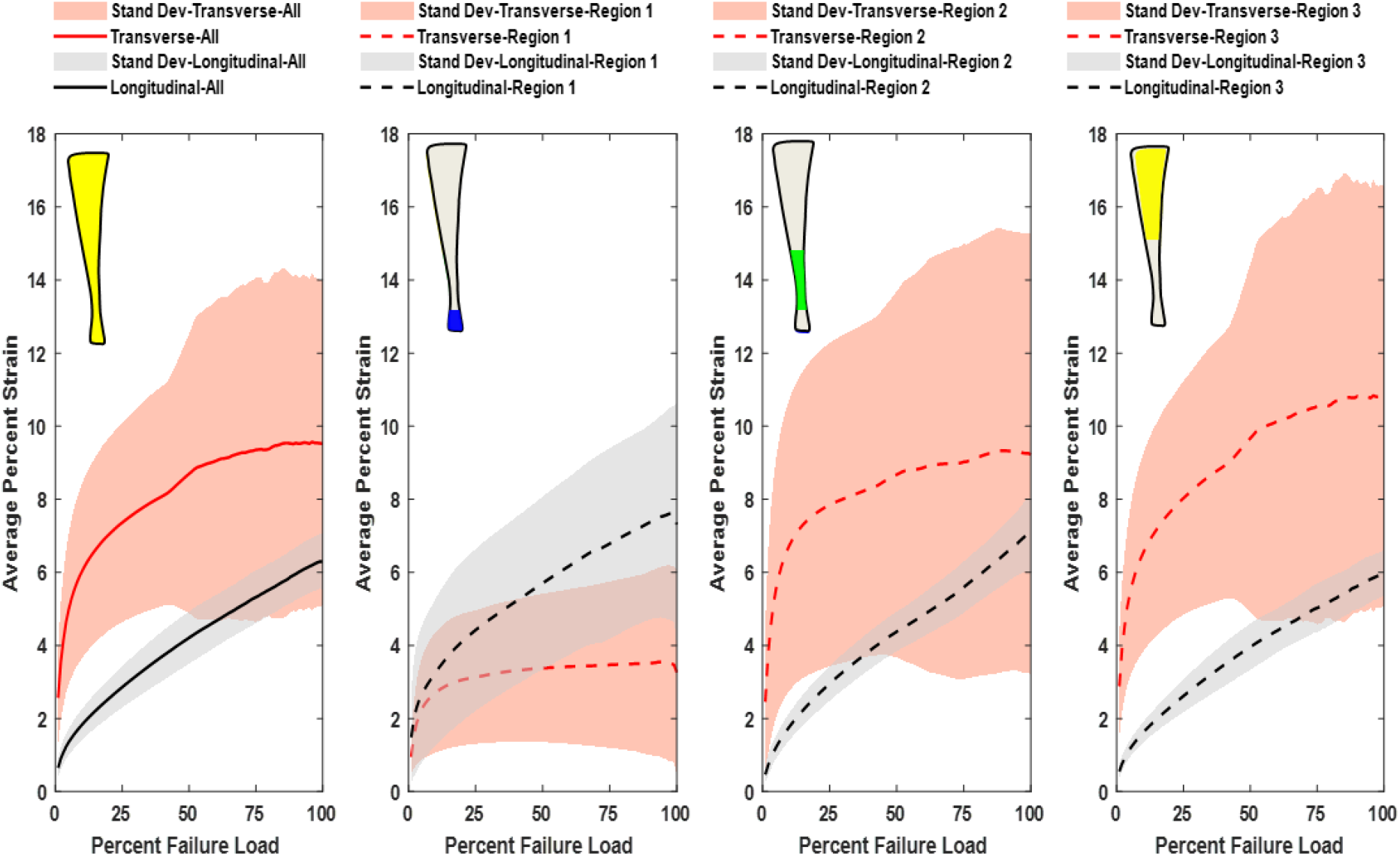
Average percent transverse and longitudinal strain as a function of percent failure load for the entire surface (A) and regions 1 (B), regions 2 (C), and regions 3 (D). Under increasing load, regions 2 and 3 experience more transverse strain than longitudinal strain, which indicates that the samples are widening more than elongating under tensile stress. Region 1 (B) experienced more longitudinal strain than transverse strain throughout loading.

### Poisson’s Ratio as a Function of Loading

Average Poisson’s ratio of the entire surface remained negative at all points during testing to failure (Fig. 8A). Poisson’s ratios were at their most negative under low loads, reaching below - 6 in region 3 (Fig. 8C). Poisson’s ratios increased as loading continued and plateaued as the failure load approached without turning positive at any time. This was a highly consistent observation regardless of failure site (Fig. 8 B).

**Figure 8:**
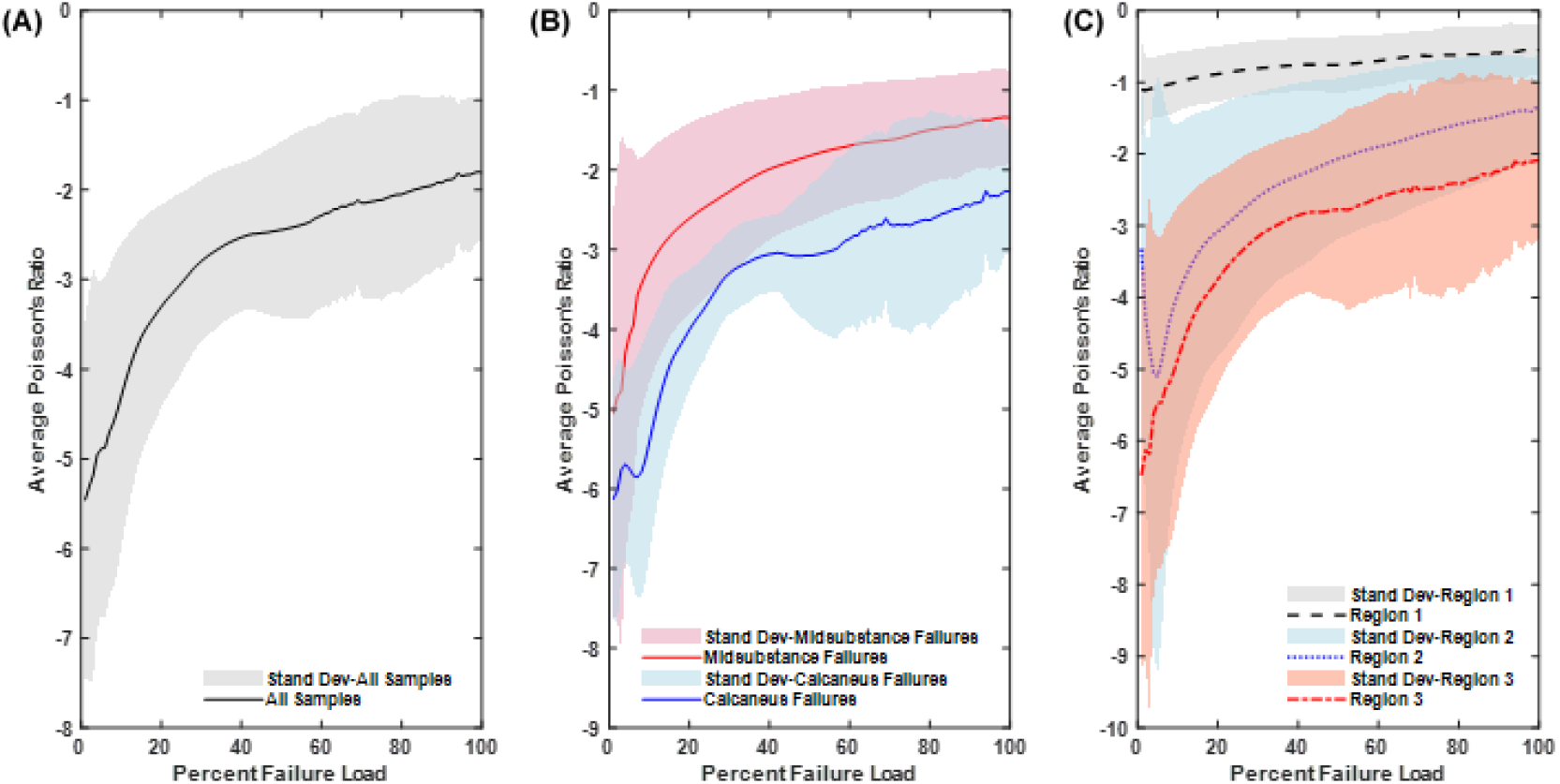
Average Poisson’s Ratio graphed as a function of percent failure load for all samples (A), by the two failure types, mid-substance tears and calcaneus avulsion (B), and by the different pre-defined regions (C). Auxetic behavior was demonstrated by the tendons regardless of failure type or region.

## Discussion

Materials with a negative Poisson’s ratio are called auxetic, a word introduced by Evans *et al*. in 1991 (Evans and Nkansah, 1991). Auxetic materials have the counter-intuitive property of becoming wider as they are stretched longitudinally (reviewed in reference (Greaves *et al*., 2011)). They have high energy absorption and fracture resistance, properties that are essential for the Achilles tendon to perform its physiological function. Various materials, such as metals, foams, zeolites, silicates, and graphene have auxetic properties, but there has been little exploration of this phenomenon in human tissues.

Nearly all previous studies of the mechanical behavior of tendons report positive Poisson’s ratios, although the range of values is high (Cheng and Screen, 2007; Chernak and Thelen, 2012; Lynch *et al*., 2003; Reese and Weiss, 2013; Vergari *et al*., 2011). However, an indication of possible auxetic behavior was published by Gatt *et al*. (Gatt *et al*., 2015) who studied flexor tendons from pigs and sheep, as well as *peroneus brevis* and Achilles tendons from two human donors. Their laboratory used traditional tensile testing methods in combination with a camera video-extensometer that tracked markings on the tendons’ surfaces. Poisson’s ratios of -1.44 and -0.39 were noted for human Achilles tendon, but this technology is outdated, less reliable and does not permit study of regional variations. Moreover, each tendon was only loaded to 2% axial strain, possibly for the practical and technical reasons alluded to in the introduction to the present paper.

The possible contribution of experimental artifacts to *ex vivo* measurement of Poisson’s ratios in tendons was subject to experimental and numerical modelling by Carniel *et al*. (Carniel *et al*., 2019), whose data highlight a large influence of clamping on specimen kinematics, even at considerable distances from the clamps. Our study did not directly clamp the tendon; the human Achilles tendons were secured by their natural insertion sites in the calcaneus and muscle. Nevertheless, cryo-clamping the distal third of the gastrocnemius muscles and the myotendinous junctions of the tendons could contribute to the auxetic behavior we observed. Specifically, compressing the muscle and tendinous junction could splay the tendon fibers and cause them to widen under tension. Additionally, because the muscle is unable to dissipate forces and absorb load, this creates an artificial boundary. It is possible that as the gastrocnemius and soleus muscle bear load, they will contract laterally, which would reduce the lateral movement of the myotendinous junction. The geometry of the tendon which narrows in the midsubstance creating wider insertions into the proximal muscle and distal calcaneus may also contribute to this mechanical response, because once stretched the tendon fibers would strain laterally to straighten. Nevertheless, MRI data from two individuals in the study of Gatt *et al*. (Gatt *et al*., 2015) indicate Achilles tendon auxeticity *in vivo*.

The values we determined for the strength of the Achilles tendon are at the high end of those reported the literature [9], consistent with the absence of pre-existing tendiopathy in our samples as determined by careful gross inspection and histology.

Luyckx *et al*. (Luyckx *et al*., 2014) were the first to apply DIC to the human Achilles tendon. Six tendons were loaded to 628.3N which, based upon our data, is only approximately 11% of failure load. Similar to the current findings, they noted considerable regional inhomogeneity and also reported significant transverse strain. However, the concept of auxeticity was not discussed.

The degree of auxetic behavior measured in the present study is remarkable both in its size and anisotropic distribution. Importantly, our findings provide a possible novel explanation for the remarkable tensile strength required to rupture the Achilles tendon during in vitro mechanical testing. Our DIC data show that the high strains imposed by loading are almost entirely absorbed by the proximal region (region 3, figure 1) in the lateral direction, thereby shielding the mid-substance where the tendon is thinnest and most prone to rupture. Its ability to do so is exhausted at 99% of the ultimate load of failure, at which point high longitudinal strain rapidly appears for the first time in the mid-substance, which quickly snaps (see video; supplemental data).

These new insights into the material basis of tendon failure, despite the limitations of testing, may have clinical relevance in the design of prevention and rehabilitation strategies. Failure testing was done on a group of relatively young, healthy Achilles tendons which, according to epidemiological studies, are less likely to sustain tendon ruptures. Auxetic behavior may be a mechanical response that allows younger tendons to dissipate large loads and forces during activity. Older individuals and people with tendinopathy are significantly more likely to sustain a tendon rupture; further research is needed to understand how aging and tendinopathy affect this behavior.

One limitation of the present study and analysis is that strain and deformation in the anterior-posterior direction were not directly evaluated. While the DIC system used is 3D, its in-plane measurements are not sufficiently robust to compute strain directly in the anterior-posterior direction. Qualitative evaluation of changes in the contour of the posterior surface of the tendon during tensile loading indicated that in the anterior-posterior direction, the tendon narrowed and hence did not exhibit auxetic behavior in that direction. However, this does not discount the auxetic behavior observed in the medio-lateral direction. In addition, it remains unclear how best to describe a material that demonstrates auxetic behavior in an asymmetric manner across different dimensions. Finally, the contributions of calf muscles to this phenomenon are unknown.

Further research is necessary to determine the molecular or structural basis for the auxetic behavior of the Achilles tendon. In vivo evidence from Gatt et al^21^ indicates auxetic or widening behavior of the tendon, but it is not clear to what extent geometry or some intrinsic property of the tendon contributes to this behavior. Studies that have tensile tested the midsubstance of the Achilles tendon, reducing the influence of the tendon’s geometry, have still noted significant transverse strains of the tendon.(Luyckx *et al*., 2014) This could be an artifact of testing but other tendons also demonstrate lateral deformation.

Tendon has few cells and type I collagen makes up over 90% of the extracellular matrix [5]. This provides considerable tensile strength in the longitudinal direction but other components, possibly including fluid flow, presumably control transverse strain. The fact that the portion of the tendon closest to the calcaneus (region 1), which displayed more longitudinal than transverse strain throughout loading, is calcified may provide a clue. An alternative explanation may reside with the twist of the collagen fascicles that exists within the human Achilles tendon. This could produce auxetic behavior of the type seen with the synthetic helical auxetic yarn of Sloan *et al* (Sloan *et al*., 2011). This will be explored in further studies; whether and how these structures are altered in tendinopathies may provide critical information.

## Acknowledgements

Authors would like to acknowledge support from the Rehabilitation Medicine Research Center, the Biomechanics Core at Mayo Clinic and the John and Posy Krehbiel Professorship in Orthopedics. Support for CN was received from the National Institute of Arthritis and Musculoskeletal and Skin Diseases grant T32AR56950. Dr. Kai-Nan An and Andrew Thoreson are thanked for critical discussion of the data.

## Legend, Supplemental Movie

Real-time, deformation measurement of major strain (A), major strain direction (B), transverse strain (C) and longitudinal strain (D) as a human Achilles tendon is loaded to failure. The heat map scale on the right indicates technical strain from -5% (blue) to 25% (red). As load is applied, strain initially accumulates in the widest region of the tendon (region 3; figure 1) in a transverse direction, with relatively little or no strain in the mid-substance (region 2; figure 1) where the tendon will eventually fail. As the tendon nears failure, the increasing transverse strain in the proximal region plateaus leading to a rapid increase in longitudinal strain in the tendon mid-substance which immediately ruptures.

## References

Carniel TA, Formenton ABK, Klahr B, Vassoler JM, de Mello Roesler CR, Fancello EA (2019) An experimental and numerical study on the transverse deformations in tensile test of tendons. J Biomech 87: 120–126.

Cheng VWT, Screen HR (2007) The micro-structural strain response of tendon. J Mater Sci 42: 8957–8965.

Chernak LA, Thelen DG (2012) Tendon motion and strain patterns evaluated with two-dimensional ultrasound elastography. J Biomech 45: 2618–2623.

Del Buono A, Chan O, Maffulli N (2013) Achilles tendon: functional anatomy and novel emerging models of imaging classification. Int Orthop 37: 715–721.

Docheva D, Muller SA, Majewski M, Evans CH (2015) Biologics for tendon repair. Adv Drug Deliv Rev 84: 222–239.

Evans KE, Nkansah MA (1991) Molecular network design. Nature 353: 124.

Ganestam A, Kallemose T, Troelsen A, Barfod KW (2016) Increasing incidence of acute Achilles tendon rupture and a noticeable decline in surgical treatment from 1994 to 2013. A nationwide registry study of 33,160 patients. Knee Surg Sports Traumatol Arthrosc 24: 3730–3737.

Gatt R, Vella Wood M, Gatt A, Zarb F, Formosa C, Azzopardi KM, Casha A, Agius TP, Schembri-Wismayer P, Attard L, Chockalingam N, Grima JN (2015) Negative Poisson’s ratios in tendons: An unexpected mechanical response. Acta Biomater 24: 201–208.

Greaves GN, Greer AL, Lakes RS, Rouxel T (2011) Poisson’s ratio and modern materials. Nat Mater 10: 823–837.

Hopkins C, Fu SC, Chua E, Hu X, Rolf C, Mattila VM, Qin L, Yung PS, Chan KM (2016) Critical review on the socio-economic impact of tendinopathy. Asia Pac J Sports Med Arthrosc Rehabil Technol 4: 9–20.

Komi PV, Fukashiro S, Jarvinen M (1992) Biomechanical loading of Achilles tendon during normal locomotion. Clin Sports Med 11: 521–531.

Lantto I, Heikkinen J, Flinkkila T, Ohtonen P, Leppilahti J (2015) Epidemiology of Achilles tendon ruptures: increasing incidence over a 33-year period. Scand J Med Sci Sports 25: e133–138.

Lewis G, Shaw KM (1997) Tensile properties of human tendo Achillis: effect of donor age and strain rate. J Foot Ankle Surg 36: 435–445.

Louis-Ugbo J, Leeson B, Hutton WC (2004) Tensile properties of fresh human calcaneal (Achilles) tendons. Clin Anat 17: 30–35.

Luyckx T, Verstraete M, De Roo K, De Waele W, Bellemans J, Victor J (2014) Digital image correlation as a tool for three-dimensional strain analysis in human tendon tissue. J Exp Orthop 1: 7.

Lynch HA, Johannessen W, Wu JP, Jawa A, Elliott DM (2003) Effect of fiber orientation and strain rate on the nonlinear uniaxial tensile material properties of tendon. J Biomech Eng 125: 726–731.

Maffulli N, Longo UG, Franceschi F, Rabitti C, Denaro V (2008) Movin and Bonar scores assess the same characteristics of tendon histology. Clin Orthop Relat Res 466: 1605–1611.

Muller SA, Todorov A, Heisterbach PE, Martin I, Majewski M (2015) Tendon healing: an overview of physiology, biology, and pathology of tendon healing and systematic review of state of the art in tendon bioengineering. Knee Surg Sports Traumatol Arthrosc 23: 2097–2105.

Reese SP, Weiss JA (2013) Tendon fascicles exhibit a linear correlation between Poisson’s ratio and force during uniaxial stress relaxation. J Biomech Eng 135: 34501.

Schepsis AA, Jones H, Haas AL (2002) Achilles tendon disorders in athletes. Am J Sports Med 30: 287–305.

Sloan MR, Wright JR, Evans KE (2011) The helical auxetic yarn - a novel structure for composites and textiles: geometry, manufacture and mechanical properties. Mechanics of Materials 43: 476–486.

Vergari C, Pourcelot P, Holden L, Ravary-Plumioen B, Gerard G, Laugier P, Mitton D, Crevier-Denoix N (2011) True stress and Poisson’s ratio of tendons during loading. J Biomech 44: 719–724.

Wren TA, Lindsey DP, Beaupre GS, Carter DR (2003) Effects of creep and cyclic loading on the mechanical properties and failure of human Achilles tendons. Ann Biomed Eng 31: 710–717.

Wren TA, Yerby SA, Beaupre GS, Carter DR (2001) Mechanical properties of the human achilles tendon. Clin Biomech (Bristol, Avon) 16: 245–251.

